# Early changes in corticomotor excitability underlie proactive inhibitory control of error correction

**DOI:** 10.1101/2021.04.25.441351

**Authors:** Borja Rodríguez-Herreros, Julià L. Amengual, Jimena Lucrecia Vázquez-Anguiano, Silvio Ionta, Carlo Miniussi, Toni Cunillera

**Author notes:** ***CONFLICT OF INTEREST*** The authors declare no disclosure of financial interests and potential conflict of interest.

## Abstract

Converging evidence indicates that response inhibition may arise from the interaction of effortful proactive and reflexive reactive mechanisms. However, the distinction between the neural basis sustaining proactive and reactive inhibitory processes is still unclear. To identify reliable neural markers of proactive inhibition, we examined the behavioral and electrophysiological correlates elicited by manipulating the degree of inhibitory control in a task that involved the detection and amendment of errors. Restraining or encouraging the correction of errors did not affect the time course of the behavioral and neural correlates associated to reactive inhibition. We rather found that a bilateral and sustained decrease of corticomotor excitability was required for an effective proactive inhibitory control, whereas selective strategies were associated with defective response suppression. Our results provide behavioral and electrophysiological conclusive evidence of a comprehensive proactive inhibitory mechanism, with a distinctive underlying neural basis, governing the commission and amendment of errors. Together, these findings hint at a decisive role for changes in corticomotor excitability in determining whether an action will be successfully suppressed.

**SIGNIFICANCE STATEMENT:** Response inhibition is a fundamental brain function that must be flexible enough to incorporate volitional goal-directed demands, along with rapid, automatic and well consolidated behaviors. Previous studies reflect a lack of consensus regarding the neural correlates subserving these two –proactive and reactive– distinct modes of inhibitory control. We combined electrophysiological recordings with behavioral measures within a paradigm of detection and correction of errors under two degrees of inhibitory control to identify genuine neural markers of proactive inhibitory control. We found evidence supporting a sustained and global –not selective– reduction of corticomotor excitability subserving successful proactive inhibition of motor responses. Our findings favor a distinctive mechanism of comprehensive inhibitory control to amend errors under a high degree of response competition.

## INTRODUCTION

Human adaptive behavior owes as much to taking suitable actions as to suppress inappropriate responses. The latter –inhibitory– processes involve both “reactive” and “proactive” control mechanisms (Aron, 2011). Reactive inhibition (e.g., pedestrians stopping at a red light) is associated to external stimuli and elicits automatic responses through habituation. Differently, proactive inhibition (e.g., halt the desire to eat in order to meet weight management goals) actively maintains abstract goal-related cues and operates with endogenous preparatory mechanisms (Miller and Cohen, 2001; Jahanshahi et al., 2015). Clear distinction between the underlying neural underpinnings sustaining proactive and reactive inhibitory processes has proven difficult due to limited behavioral paradigms, not ideally suited to isolate one from the other (Mostofsky and Simmonds, 2008; Meyer and Bucci, 2016). The present study sought to experimentally manipulate the degree of inhibitory control within a paradigm of constrained correction of errors to investigate putatively distinct neural markers of proactive inhibition with event-related potentials (ERPs) and brain oscillations.

There is cumulative evidence challenging response inhibition as a classically considered ‘unitary’ brain function (Aron et al., 2014; Raud et al., 2020). Broadly, having foreknowledge of which response must be suppressed has been associated with selective but also slower effortful proactive inhibitory mechanisms, whereas a reflexive reactive global process prevails when the priority is to stop quickly (Aron and Verbruggen, 2008; Greenhouse et al., 2012). The dissociation of these two mechanisms of response inhibition resembles the central hypothesis of the dual mechanisms of control (DMC) framework, which postulates that cognitive control operates via two distinct control modes with different temporal dynamics (Braver, 2012). Specifically, DMC account posits that variability in cognitive control arises from a transient activation of the lateral prefrontal cortex (PFC) when reactive control acts as a ‘late-correction’ mechanism depending on the detection of cognitively demanding events, and a sustained activation of lateral PFC when proactive control relies upon their anticipation and prevention. Nevertheless, neuroimaging studies have failed to dissect two separate fronto-striatal brain networks for reactive and proactive inhibitory processes, which exhibit substantial overlapping of activated areas including the right inferior frontal cortex (rIFC), the striatum and the subthalamic nucleus (Aron and Poldrack, 2006; Chikazoe et al., 2009; Majid et al., 2013; Cunillera et al., 2014).

Inspired by previous electrophysiological evidence showing different inhibitory mechanisms when stopping or changing a planned response (De Jong et al., 1995; Kramer et al., 2011), we manipulated the degree of inhibitory control by either forbidding or encouraging participants to correct their erroneous responses. We used a ‘stop-change paradigm’ in which a switch of the direction of the stimulus elicited not only the suppression but also the immediate execution of a fixed alternative response. Under two different degrees of inhibitory control, this behavioral paradigm allowed us to create situations of high attentional loading –in which proactive inhibition is at use–, to cope with overcorrection of errors. We investigated the time-course of error-and response-related ERPs as well as two candidate ERPs for successful response inhibition, the N2 and the P3 components. The cascade of electrophysiological correlates associated with the response preparation and execution was also assessed by the study of the oscillatory theta-and beta-frequency bands. Under a high degree of response competition, we expected the forbidding and the encouraging of error corrections to elicit reliable neural signatures of proactive inhibitory control, which would be concurrently associated with successful response suppression.

## MATERIALS AND METHODS

### Participants

Nineteen healthy subjects (12 women; mean age = 20.7 years, SD = 2.46 years) participated in the experiment. All the participants had normal or corrected-to-normal visual acuity and no history of neurological or psychiatric disorders. All participants were naïve with respect to the experimental procedures and the hypothesis of the study. Prior to their inclusion in the study, participants provided written informed consent. The study was approved by the local Ethics Committee according to the Declaration of Helsinki. All participants were right-handed, as assessed by the Edinburgh handedness questionnaire (Oldfield, 1971), and were paid or received extra course credits after completing the experiment.

### Stimuli and procedure

Participants sat in front of a table positioned 45-50 cm below their eyes. Visual stimuli were presented using Presentation^®^ software (v.0.52, Neurobehavioral Systems, Inc., Berkeley, CA, https://www.neurobs.com), running on Windows XP-32SP3 in an Intel Core-i3 computer, and displayed on a 21” Philips Brilliance 202P4 CRT monitor with a refresh rate of 144 Hz and a resolution of 1024 x 768 pixels. We employed a modified version of the Eriksen flanker task (Eriksen and Eriksen, 1974). A horizontal array of five green arrows (4×10 cm) was presented for 400 ms in the center of the screen at a Euclidian distance of 60 cm and with a visual angle of ∼3.4°. The stimulus onset asynchrony (SOA) was randomly established between 1000 and 1200 ms. Participants were instructed to respond to the direction of the central arrow by using the right index finger for right-directed responses and the left index finger when the arrow pointed leftwards. The four surrounding arrows were either congruent (*Compatible*) or incongruent (*Incompatible*) with respect to the direction of the central arrow. The proportion of compatible and incompatible trials was set to 40/60%.

The novel and relevant aspect that differentiates our task from the classical Eriksen version is that a switch in the direction of the central arrow –equally distributed between compatible and incompatible conditions– was introduced in 25% of trials (**Figure 1**). In these switch trials, participants had to avoid responding to the initial direction of the central arrow and respond instead in accordance with its direction after the switch. The onset of the switch (hereafter switch-signal delay, *SwSD*), defined as the time between the go stimulus and the switch signal, was initially set to 200 ms and adapted dynamically on a trial-by-trial basis by means of a staircase-tracking algorithm to compensate for differences between participants (Osman et al., 1986; Logan et al., 1997; Band and van Boxtel, 1999). The step size, that is, the difference in the SwSD from one switch trial to the other was ±20 ms. Put simply, after a correct response in a switch trial, the SwSD was increased by 20 ms. In contrast, the SwSD was reduced by 20 ms after an error, thereby modulating, respectively, the difficulty in the next switch trial. Dynamic tracking procedure ensured an overall ratio of p(*response*|*switch*) of 0.5 in each participant, regardless of their baseline performance.

**Fig. 1.**
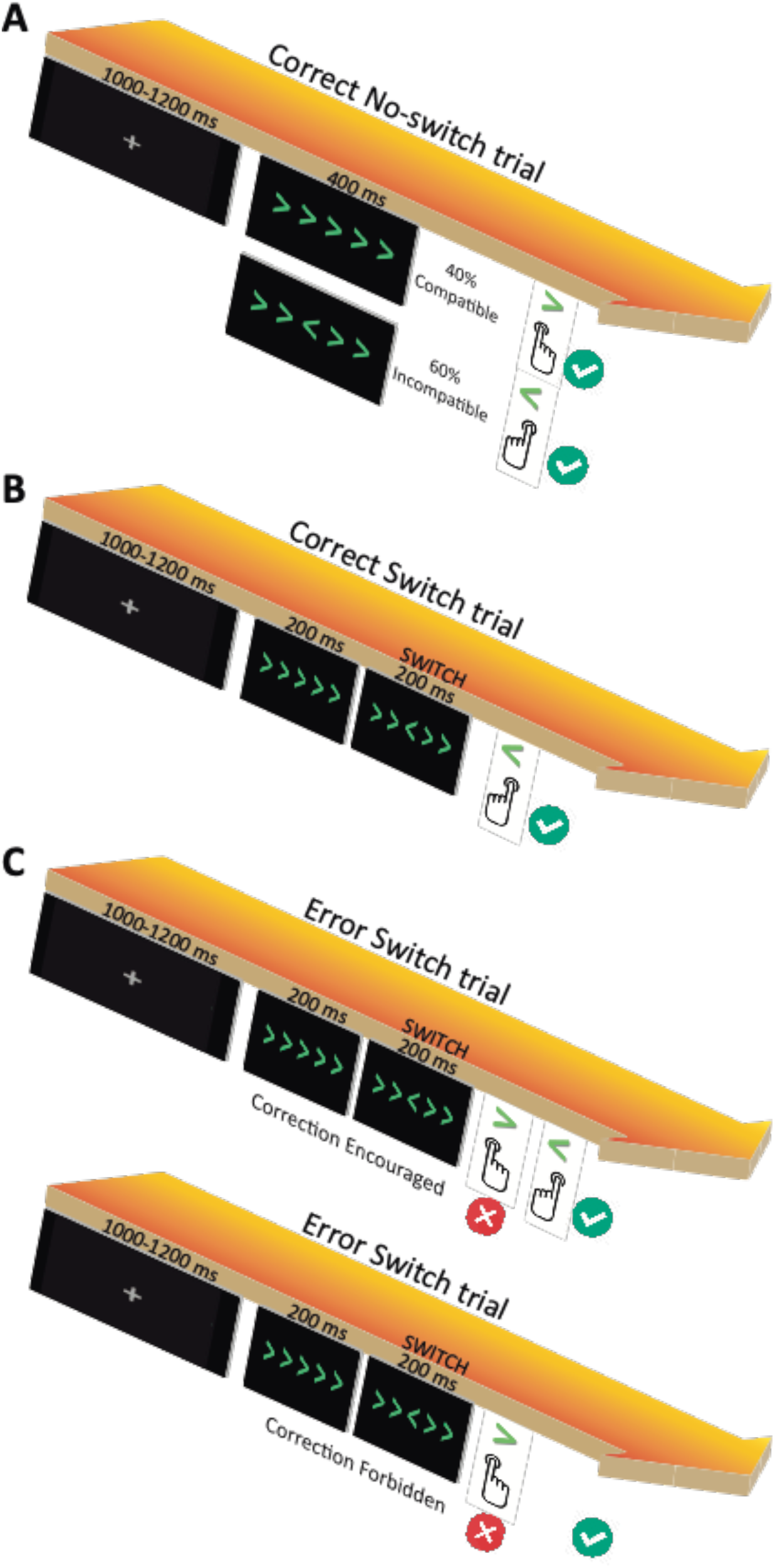
**(A)** Examples of correct responses to compatible and incompatible no-switch trials. **(B)** Example of a correct response to a switch trial. **(C)** Examples of erroneous responses to a switch trial both in the Correction Encouraged (error must be amended) and the Correction Forbidden (error must not be amended) conditions. The switch is presented in 25% of trials.

### Experimental design

The experiment consisted of two sessions. The first session was designed as a training block of 15 min to practice the standard flanker task without switch trials (i.e., participants were instructed to respond to the direction of the central arrow with the corresponding index finger). The second session corresponded to the real experiment with the modified ‘flanker-switch’ paradigm. Participants performed this second session divided in two blocks under two different sets of instructions. In one block, participants were asked to correct their responses in case they committed an error (*Correction Encouraged* condition); whereas in the other block, participants were instructed to inhibit any corrective counteracting response after an error (*Correction Forbidden* condition). The order of the two blocks was counterbalanced among participants. Each experimental block consisted of six consecutive runs of 240 trials each, resulting in a total of 2880 trials for the whole session. Before starting a block, participants performed 20 practice trials to get familiar with the task. The total duration of the experiment was approximately 3 hours.

### Behavioral data analysis

We registered the response in each trial and measured the response time (RT). In no-switch trials, RT was the time between the trial onset (i.e., the display of the arrows) and the finger tapping, whereas in switch trials RT was defined as the time between the switch onset and the response. A two-sided Wilcoxon matched-pairs test was performed to compare the percentage of correct responses, both in switch and no-switch trials, for each correction instruction. The evolution across time of the SwSD was used as an indicator of the inhibitory performance (i.e., SwSD would increase with successful response inhibition and vice versa). To measure the latency of the inhibitory process, we computed the switch-signal reaction time (SwSRT) following the integration method (Logan, 1981; Logan and Cowan, 1984). SwSRT was estimated by subtracting the averaged SwSD from the *n*^th^ RT separately for each condition, where *n* is obtained by multiplying the number of no-switch RTs in the distribution by the overall p(respond/stop-signal). We conducted a 2 x 2 repeated measures analysis of variance (ANOVA) with factors Correction instruction (Encouraged, Forbidden) and Congruency (Compatible, Incompatible) to determine the influence of the two degrees of inhibitory control on the compatibility effect. We also compared RTs between erroneous and correct responses in a 3 x 2 repeated measures ANOVA with factors Correction instruction (Encouraged, Forbidden), Switch (No-Switch trial, Switch trial) and Performance (Correct, Error). The SwSD, the SwSRT, as well as the RT differences between correct and error switch responses following a previous switch error, were entered into separate 2 x 2 ANOVAs with Correction instruction (Encouraged, Forbidden) and Performance (Correct, Error) as fixed factors. The error rate in switch trials after a previous switch error was also compared between Encouraged and Forbidden correction conditions.

### Electrophysiological recording and analysis

The electroencephalogram (EEG) was recorded with a BrainAmp DC amplifier (*Brain Products GmbH*) from 28 scalp electrodes mounted in an elastic cap (*EasyCap*) and displayed in accordance with the standard 10/20 system. The electrodes were located at Fp1/2, F3/4, F7/8, FC1/2, FC5/6, T3/4, C3/4, CP5/6, CP1/2, T5/6, P3/4, O1/2, AFz, Fz, Cz, and Pz). The EEG signal was sampled at 250 Hz, referenced online against the right mastoid electrode and re-referenced offline against the half mean of the left mastoid. Electrode impedances were kept below 5 kΩ during the experiment. Eye movements were monitored and recorded by an electrode situated below the right eye. Muscular artifacts and eye blinks were removed offline, first using a voltage threshold of ±100 μV and then detecting abrupt voltage changes of ±25 μV within time windows of 10 ms. Stimulus-locked (S-Locked) and response-locked (R-Locked) epochs were computed separately from the resulting artifact-free signal for each condition. The length of the epochs was 1000 ms for both modalities (−100 to 996 ms in S-Locked ERPs, and -400 to 600 ms in R-Locked ERPs). Before averaging, the EEG signal was filtered offline with a high-pass filter of 0.1 Hz, to remove possible electrode drifts. Subsequently, ERPs epochs were filtered with 12 Hz and 20 Hz low-pass filters for R-locked and S-locked epochs, respectively, but only for presentation purposes. We characterized the error-related ERPs (ERN-Pe compound) and two candidate ERPs commonly reported for successful response inhibition: the frontolateral N2 wave and the frontocentral P3 components. Mean amplitude measures in switch trials were obtained and entered into a 2 x 2 x 4 repeated-measures ANOVA, with Correction instruction (Encouraged, Forbidden), Performance (Correct, Error) and the four midline electrode locations (AFz, Fz, Cz and Pz) as the studied factors.

In addition to ERPs, Time-Frequency (TF) analysis was performed by convoluting single-trial data with a complex Morlet wavelet:

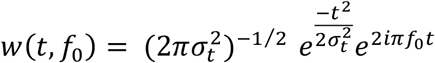

where the relation f_0_/σ_*f*_ (where σ_*f*_ = 1/(2πσ_*t*_)) was set to 6.7 (Tallon-Baudry et al., 1997). The frequencies studied ranged from 1 to 40 Hz, with a linear increase of 1 Hz. The time-varying energy –defined as the square of the convolution between wavelet and signal– was computed for each trial and averaged separately for each participant. TF contents were averaged switch-signal locked epochs. Mann-Wilcoxon sum tests were performed for all frequencies and time points to test for significant differences in power between the two degrees of inhibitory control. Percentage of increase/decrease in power for these conditions were entered into the analyses of variance with Correction instruction (Encouraged, Forbidden) and Electrode (AFz, Fz, Cz, Pz) as factors. Significance threshold was set at p<0.01 and only significant clusters larger than 100 ms were considered. The Greenhouse-Geisser epsilon correction was applied when necessary in all ERP and TF analyses (Jennings and Wood, 1976).

### Lateralized readiness potential (LRP) and current source density analysis (CSD)

As an index of prepotent motor activity, we measured LRPs to quantify the motor preparatory activity elicited with the two correction instructions. LRPs were assessed using C3 and C4 electrodes, in which the amplitude of the readiness potential is maximum (Kutas and Donchin, 1980). The LRP is computed by a double subtraction as shown in the following equation: LRP = left hand (C4–C3) – right hand (C4– C3). Left and right hands refer to the expected correct hand, and (C4–C3) is the difference in electrical potential between these electrodes (Gratton et al., 1988; Smid et al., 1992). The resulting LRP component is negative if participants produce correct responses and positive when they produce a response with the alternative hand, such as when correcting an error. For statistical analysis, mean amplitude values in a 100 (±50) ms time window around the peak of the motor preparatory activity –located in the grand average waveform separately for correct and error switch trials– were introduced in a 2 x 2 repeated-measures ANOVA with Correction instruction (Encouraged, Forbidden) and Performance (Correct, Error) as the studied factors. All LRP data was filtered with 12 Hz low-pass filter for the statistical analysis. Huynh-Feldt epsilon correction was applied when necessary.

We used current source density (CSD) –a reference-free technique that computes the second spatial derivative (Laplacian) of the scalp electric potential–, to obtain a reliable and more precise measure of the inhibitory activity in the motor cortex. Laplacian removes the noncortical-induced volume conduction to improve the spatial resolution and provides the location, direction and intensity of the radial current flow that determines an ERP topography (Mitzdorf, 1985; Perrin et al., 1989). We obtained surface laplacian computed on individual ERP data from C3 and C4 electrodes –positioned bilaterally over the motor cortex–, using the MATLAB-based CSD toolbox (Kayser, 2009) with EEGLAB (Delorme and Makeig, 2004). We therefore transformed all the averaged ERP waveforms into reference-free CSD estimates (μV/cm^2^ units, head radius = 10 cm). The interpolation was computed using the spherical spline surface Laplacian with computation parameters (50 iterations; spline flexibility *m* = 4; smoothing constant λ = 10^−5^) previously established for our 28-channel recording montage.

We collapsed CSD estimates across all conditions, but separately with respect to hemispheric laterality to create contralateral and ipsilateral responses. In other words, the contralateral condition averaged CSD estimates at C3 from right-hand responses with CSD estimates at C4 from left-hand responses, and vice versa for the ipsilateral condition. We filtered the S-locked CSD estimates with a second order Infinite Impulse Response (IIR) Butterworth low-pass filter (12-40 dB) at 20 Hz and entered them into a 2 x 2 repeated-measures ANOVA with Correction instruction (Encouraged, Forbidden) and Performance (Correct, Error) as factors. Based on previous studies on inhibitory control (Burle et al., 2016), we measured the area under the CSD waveform in a predefined time window as well as the slope of the CSD deflection in that window to obtain a second measure not affected by the selected baseline. The slopes were computed by fitting a linear regression to the signal in the time window of interest. Huynh-Feldt and Greenhouse-Geisser epsilon corrections were applied when necessary.

## RESULTS

### Behavioral results

Participants successfully inhibited the correction in approximately half of the switch trials when correction was forbidden (50.8 ± 3%), indicating an optimal implementation of the staircase-tracking algorithm. When the correction was encouraged, participants were also able to correct their erroneous responses in comparable proportions (48.7 ± 4.4%, *p* = 0.11). Accuracy in no-switch trials reached similar levels (*p* = 0.52) with both the Encouraged (94.6 ± 4.3%) and the Forbidden (92.9 ± 5.9%) correction instructions.

Figure 2 and **Table 1** show the main behavioral results. We reproduced a compatibility effect both when encouraging and forbidding error correction (Compatibility: *F*_1,18_ = 156.1, *p* = 2.6e-10, *η*_*p*_^2^ = 0.9; **Figure 2A**). RTs in incompatible trials were 21 ms (*t*_(18)_ = -7.9, *p* = 1.8e-9, *d* = 1.28) and 20 ms (*t*_(18)_ = -6.58, *p* = 1e-7, *d* = 1.07) longer when the correction was encouraged and forbidden, respectively. **Figure 2B** displays RTs in switch and no-switch trials as a function of the performance and the correction instruction. Overall, erroneous responses (298 ms) exhibited shorter RTs than correct (375 ms) responses (Performance: *F*_1,18_ = 184.9, *p* = 6.6e-11, *η*_*p*_^2^ = 0.91). Nevertheless, we observed that this decrease in RT was four times larger in switch trials than in no-switch trials (123 vs. 31 ms; Switch x Performance: *F*_1,18_ = 36.9, *p* = 9.7e-6, *η*_*p*_^2^ = 0.67). The SwSRT (273 vs. 272 ms) and the Go RT (376 vs. 371 ms) were practically identical in the Encouraged and the Forbidden correction conditions (*p* > 0.28 for all comparisons, **Figure 2C**). The progression of the SwSD across time for each correction instruction is displayed in **Figure 2D**. When error correction was encouraged, we observed a decrease of the SwSD in the first trials down to an average slightly below 100 ms, remaining consistently at this value across 200 trials. We found the same pattern when error correction was forbidden. After 200 trials, an additional decrease of the SwSD was observed in both conditions to an average of ∼75 ms.

**Table 1.** Summarized mean values for all behavioral parameters analyzed in the study. We report mean and SD separated for the Encouraged and Forbidden correction conditions.

|  | Correction<br>Encouraged | Correction<br>Forbidden | *p*-value |
| --- | --- | --- | --- |
| **No-switch trials** |  |  |  |
| *Accuracy (%)* | 94.6 (4.3) | 92.9 (5.9) | 0.52 |
| *RT* |  |  |  |
| Correct (ms) | 376.8 (72) | 373.2 (74) | 0.76 |
| Error (ms) | 320.1 (68) | 316.3 (74) | 0.75 |
| **Switch trials** |  |  |  |
| *Accuracy (%)* | 48.7 (4.4) | 50.8 (3) | 0.11 |
| *RT* |  |  |  |
| Correct (ms) | 373.1 (79) | 380.7 (82) | 0.86 |
| Error (ms) | 257.1 (68) | 253.7 (63) | 0.52 |
| *SwSD (ms)* | 95.3 (54) | 95.1 (57) | 0.98 |
| *SwRT (ms)* | 273.1 (19) | 272.4 (18) | 0.91 |

**Fig. 2.**
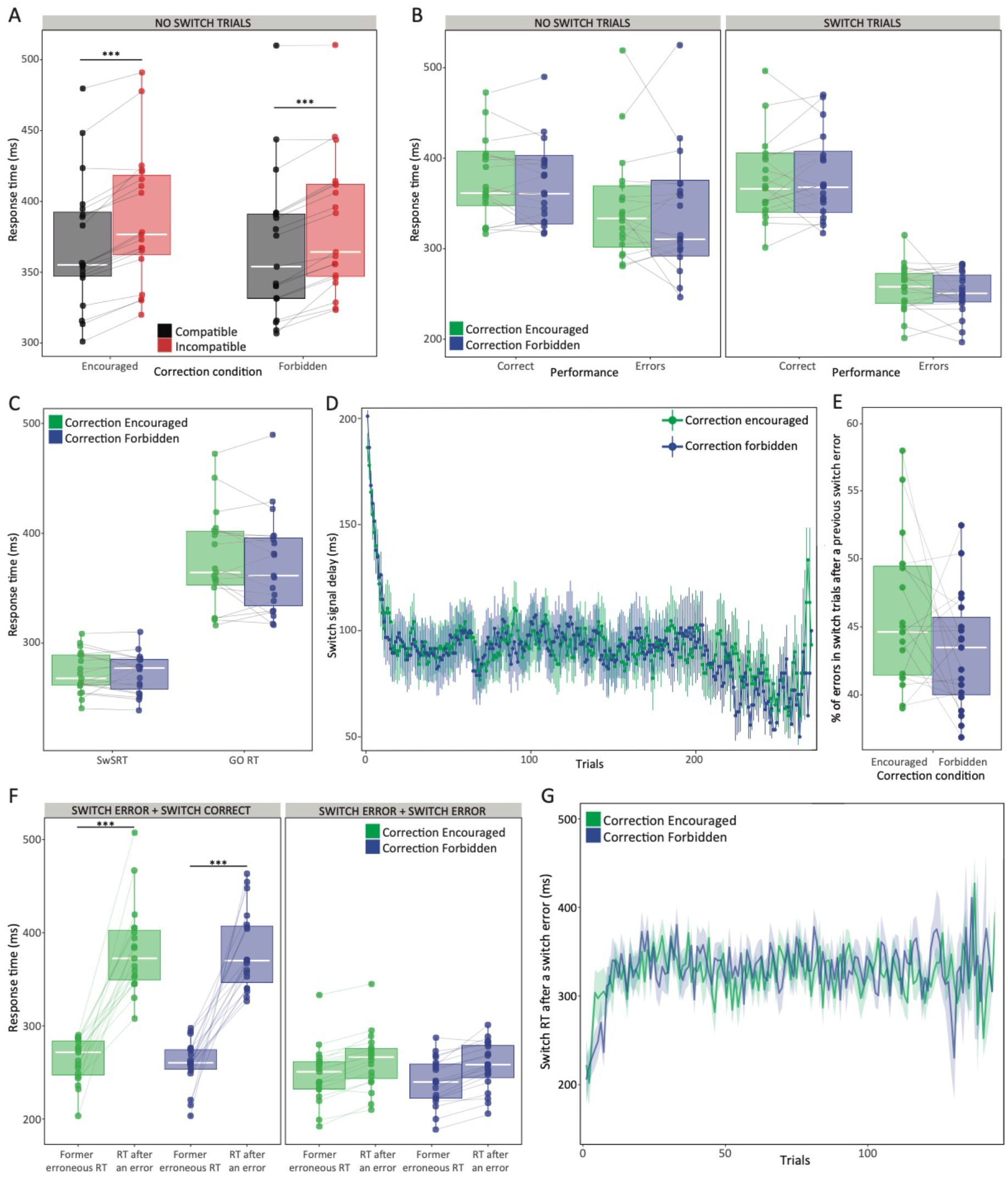
**(A)** Average RT for compatible and incompatible no-switch trials in the Encouraged and Forbidden correction condition. **(B)** Average RT for correct and erroneous responses separately for no-switch and switch trials in the two degrees of inhibitory control. **(C)** Average SwSRT and Go RT in the Encouraged and Forbidden correction condition. **(D)** Changes in the SwSD for each correction instruction are illustrated with respect to trial number. Solid thick lines connect 300 points corresponding to the inter-subject mean for each trial, while the error bars represent the standard error of the mean (SEM). **(E)** Proportion of two consecutive erroneous responses to a switch in the Encouraged and Forbidden correction condition. **(F)** Average RT for correct and erroneous responses following an error in switch trials, separately for the two correction instructions. **(G)** Changes in the RT after an error in switch trials for each correction instruction are illustrated with respect to trial number. Shaded areas represent the SEM. In all boxplots, the bold white line shows the median, and the bottom and top of the box show the 25th (quartile 1 [Q1]) and the 75th (quartile 3 [Q3]) percentile, respectively. The upper whisker ends at highest observed data value within the span from Q3 to Q3+1.5 times the interquartile range (Q3–Q1), and lower whisker ends at lowest observed data value within the span for Q1 to Q1-(1.5 * interquartile range). Points not reached by the whiskers are outliers. Significant post hoc group comparisons are represented by solid lines above. \**p*<.05; \*\**p*<.01; \*\*\**p*<.001.

We also evaluated the performance of each participant after the commission of an error in a switch trial. The error rate after a previous switch error was comparable between Correction encouraged and Correction forbidden conditions (Correction Instruction: *F*_1,18_ = 2.68, *p* = 0.12; **Figure 2E**). After a switch error, we observed that the RT of the next switch trial was on average 67 ms longer (*F*_1,18_ = 127.2, *p* = 1.4e-9, *η*_*p*_^2^ = 0.88). **Figure 2F** reveals that this RT slowing down was clearly dependent on the performance of that subsequent switch trial (*F*_1,18_ = 68.1, *p* = 1.6e-7, *η*_*p*_^2^ = 0.79): An additional erroneous response exhibited an RT only 16 ms larger than that of the former error (244 vs 260 ms, *t*_(18)_ = -2.4, *p* > 0.05 after correction for multiple comparisons); whereas a correct response to switch showed an RT 120 ms larger than that of the preceding switch error (260 vs 381 ms, *t*_(18)_ = -13.5, *p* = 7e-16, d = 2.19). These differences were indicative of a ‘switch cost’ in correct responses to switch, which was analogous when error correction was encouraged or forbidden. A trial-by-trial analysis of the switch RTs after a switch error shows that, on average, RTs increased quickly after the first trials to a value within the 300-360 ms, being consistent through blocks and between correction instructions (**Figure 2G**).

### ERP results

#### Compatibility and error-related effects

The compatibility effect was reflected on the modulation of the N2 component at the Cz electrode in no-switch trials (Compatibility: *F*_1,18_ = 4.9, *p* = 0.038, *η*_*p*_^2^ = 0.34). **Figure 3A** shows that the S-locked (*incongruent – congruent*) difference waveform was not different between the two degrees of inhibitory control (Correction Instruction: *F*_1,18_ = 3.0, *p* = 0.1; Correction Instruction x Compatibility: *F*_1,18_ = 0.06; *p* = 0.82), indicating that encouraging or forbidding the correction of errors exerted no influence in the early N2-conflict processing of congruent and incongruent sets of stimulus arrays.

**Fig. 3.**
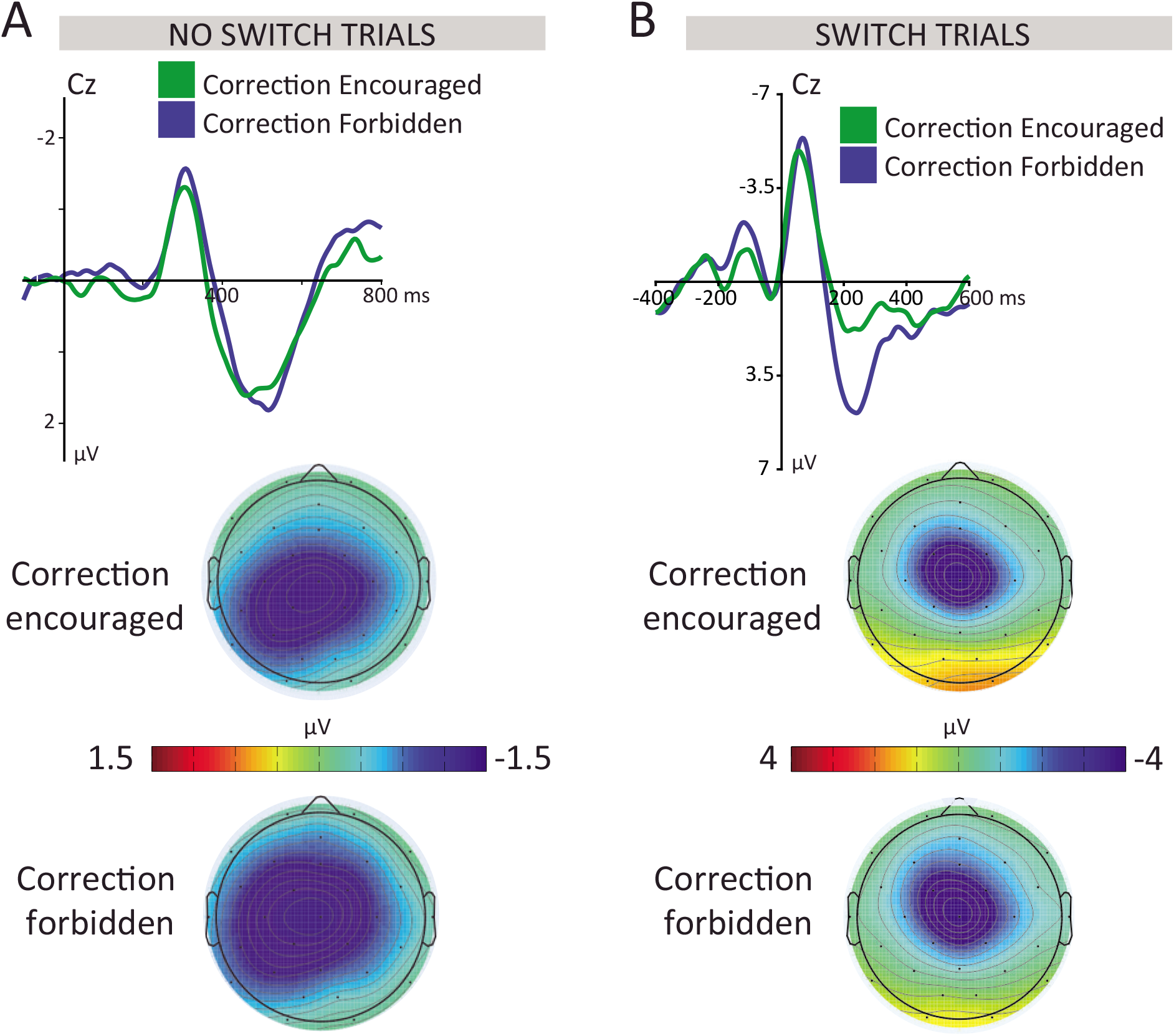
**(A)** Grand average Stimulus-locked electrophysiological correlates of the compatibility effect in no-switch trials. Incongruent minus congruent difference waveform in Cz electrode, together with the 2D isovoltage topographical mapping (283-333 ms) illustrating the scalp distribution of the N2 component for each correction condition. **(B)** Response-locked grand average and 2D isovoltage topographical mapping (45-95 ms) of the error *minus* correct difference waveform eliciting an ERN-Pe compound in switch trials for each correction condition.

The neural correlates of the errors committed in switch trials were examined by analyzing the R-locked (*error – correct*) difference waveform on the ERN-Pe compound (**Figure 3B**). While no difference in the modulation of the ERN was found when comparing the two degrees of inhibitory control (Correction Instruction: *F*_1,18_ = 2.77, *p* = 0.11; Correction Instruction x Electrode: *F*_3,54_ = 0.42, *p* = 0.61), we observed a significantly larger Pe amplitude when the correction of errors was forbidden (*F*_1,18_ = 9.31, *p* = 0.007, *η*_*p*_^2^ = 0.34).

#### Inhibitory processing: N2 and P3 components

**Figure 4A** shows the modulation of the N2 component as a function of performance and correction instructions. N2 amplitude was larger in failed compared to successful response inhibition (Performance: *F*_1,18_ = 15.88, *p* = 8.6e-4, *η*_*p*_^2^ = 0.47). Furthermore, the instructions given to the participants also affected the amplitude of the N2 (Correction instruction x Electrode: *F*_3,54_ = 16.11, *p* = 5.8e-5, *η*_*p*_^2^ = 0.47). This interaction was driven by a marginally larger N2 amplitude in frontal electrodes when corrections were encouraged (*AFz*: *t*_(18)_ = 2.04, *p* = 0.056, *d* = 0.2; *Fz*: *t*_(18)_ = 1.87, *p* = 0.08, *d* = 0.2), but a reversed pattern in posterior locations (*Pz*: *t*_(18)_ = -3.16, *p* = 0.005, *d* = -0.4), as shown in **Figure 4B**. We also found a significant interaction between the correction instruction and the performance (*F*_1,18_ = 4.77, *p* = 0.04, *η*_*p*_^2^ = 0.21). The decomposition of this interaction, however, revealed that N2 amplitude did not differed between the two correction instructions, neither in correct (*t*_(18)_ = 0.94, *p* = 0.36) nor in error switch trials (*t*_(18)_ = -0.76, *p* = 0.45).

**Fig. 4.**
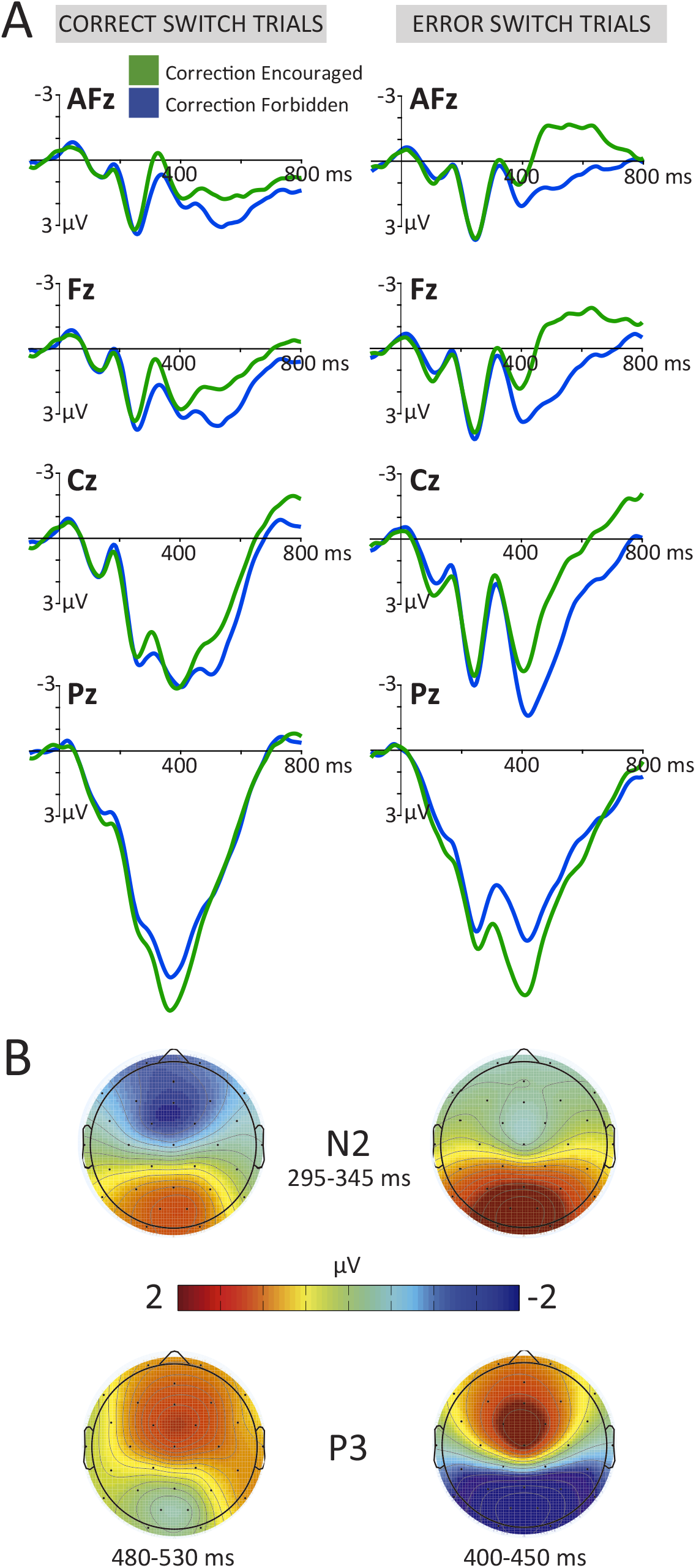
**(A)** Stimulus-locked grand average from the midline electrodes for the correct (left) and error (right) switch trials, separately for the Encouraged (green) and the Forbidden (blue) correction condition. **(B)** Topographical mapping of the N2 and the P3 components indicating the analyzed time window.

The analysis of the modulation of the P3 component revealed a different pattern when compared to the preceding N2. We found a significant Correction instruction x Electrode interaction (*F*_3,54_ = 23.96, *p* = 1.2e-5, *η*_*p*_^2^ = 0.57), driven by a significantly larger P3 amplitude in posterior regions when participants were instructed to correct their errors (*Pz*: *t*_(18)_ = -3.06, *p* = 0.007, *d* = -0.4, **Figure 4B**). The second positive phasic deflection following the P3 component –peaking at 525 ms in anterior electrodes– was also considered for the analysis. The results of the ANOVA revealed a significant main effect of correction instruction (*F*_1,18_ = 8.65, *p* = 0.009, *η*_*p*_^2^ = 0.33), and performance (*F*_1,18_ = 10.88, *p* = 0.004, *η*_*p*_^2^ = 0.38). Furthermore, we found a significant Correction instruction x Performance x Electrode triple interaction (*F*_1,18_ = 10.73, *p* < 0.001, *η*_*p*_^2^ = 0.37), suggesting that differences between erroneous and correct responses in each electrode location were modulated by the degree of inhibitory control exerted by the task demands. To further decompose the source of this interaction, we generated a ‘performance’ (*error – correct*) difference waveform separately for the Encouraged and Forbidden correction conditions. Further post-hoc comparisons showed that the ‘performance’ difference waveform differed between correction instructions only in frontal (*AFz*: *t*_(18)_ = -2.95, *p* = 0.01, *d* = -0.6; *Fz*: *t*_(18)_ = -2.88, *p* = 0.01, *d* = - 0.6), but not in medial and posterior locations (*Cz*: *t*_(18)_ = -1.74, *p* = 0.1; *Pz*: *t*_(18)_ = 1.1, *p* = 0.28).

#### Time-frequency results

Figure 5. shows the switch-locked TF and statistical maps of the event-related oscillatory activity recorded from Afz, Fz, Cz and Pz electrodes during correct responses to switch for both Encouraged and Forbidden correction conditions. **Figure 6** shows the equivalent for erroneous responses. Overall, we found an increase of the theta activity peaking around 400 ms after the switch onset in the two correction conditions. This increase was larger in central locations (t_(18)_ > 4.8, *p* < 0.001 for all comparisons). Also, mu (12-15 Hz) and beta (20-30Hz) activity were desynchronized within the 200-800 ms post-switch time interval, especially in central and posterior locations (*F*_1,18_ > 3.4, *p* < 0.05 for both correct and erroneous responses). Lastly, we observed a rebound of the beta activity between 700 ms and 1000 ms after the switch. The statistical maps across frequencies and time in correct responses did not exhibit any significant TF cluster when comparing the two degrees of inhibitory control (**Figure 5**). In contrast, we found that beta synchronization was larger between 700 ms and 900 ms after the switch onset when participants failed to withhold the error correction, prominently in anterior regions (**Figure 6**). The analysis of the latency onset determined that the beta synchronization after failing to inhibit a correction started 150 ms earlier (850 ms vs 1000 ms after the switch onset).

**Fig. 5.**
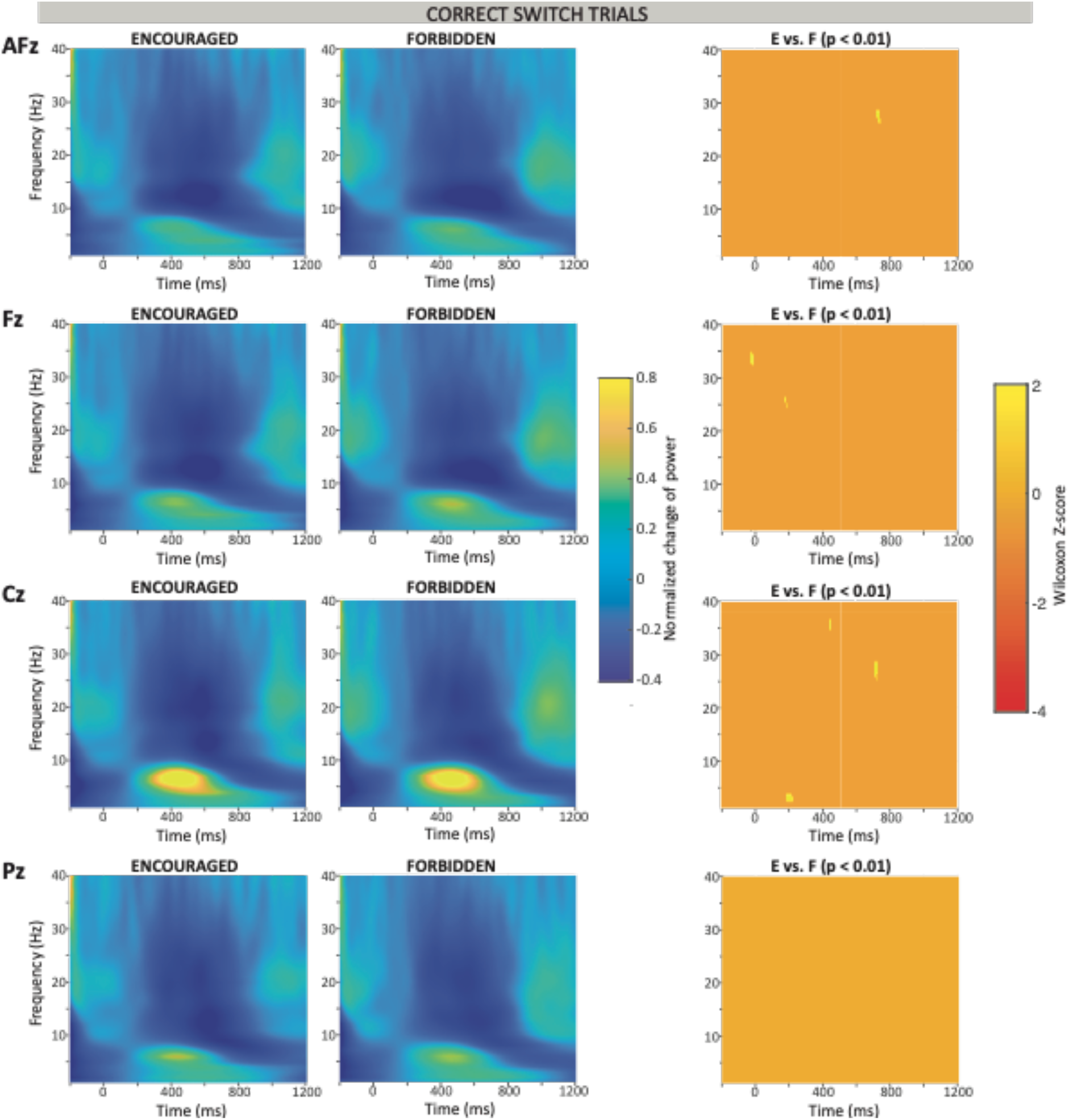
Switch-locked TF maps from frequencies between 1 to 40 Hz obtained from the midline electrode locations for correct switch trials in both Encouraged and Forbidden correction conditions. Changes of power relative to baseline (−200 to 0 ms prior to the switch). The right column presents point-by-point Mann-Wilcoxon tests between Encouraged and Forbidden correction conditions. Only significant p-values (p < .01) are represented.

**Fig. 6.**
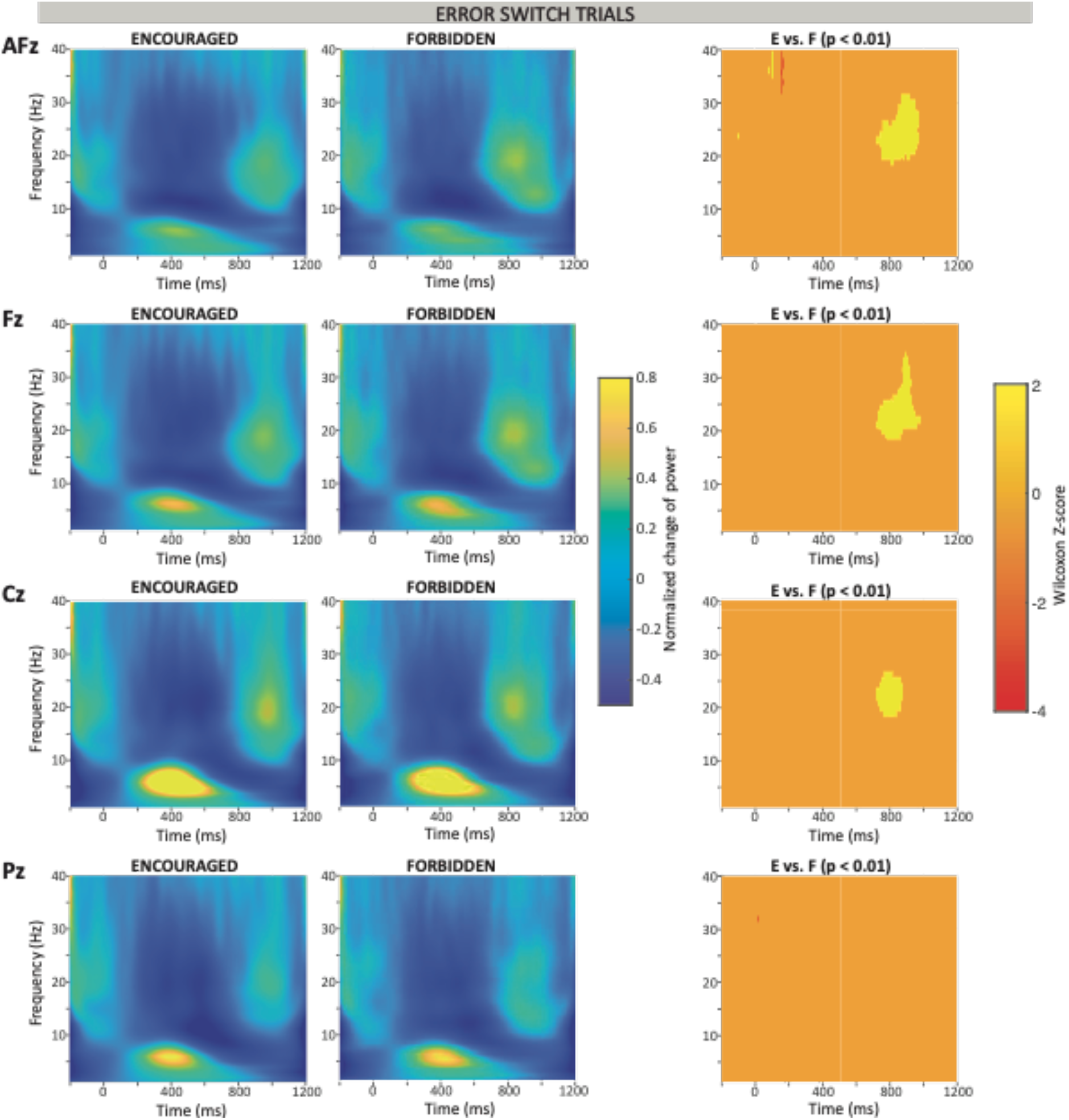
Switch-locked TF maps from frequencies between 1 to 40 Hz obtained from the midline electrode locations for error switch trials in both Encouraged and Forbidden correction conditions. Changes of power relative to baseline (−200 to 0 ms prior to the switch). The right column presents point-by-point Mann-Wilcoxon tests between Encouraged and Forbidden correction conditions. Only significant p-values (p < .01) are represented.

#### LRP and CSD results

We measured the motor preparatory activity separately in correct (1^st^ peak at 184 ms; 2^nd^ peak at 360 ms) and error switch trials (1^st^ peak at 184 ms; 2^nd^ peak at 368 ms). As expected, the LRP indicating motor preparatory activity in the first time window (134-234 ms) displayed a remarkable polarization with opposite sign associated to the performance in switch trials (*F*_1,18_ = 228.34, *p* = 1.1e-11, *η*_*p*_^2^ = 0.93; **Figure 7A**). Before a correct response to the switch, we however found a similar inhibitory pattern of the motor preparation activity when corrections were encouraged or forbidden (*F*_1,18_ = 1.47 *p* = 0.24). The interaction between the correction instruction and the performance was not significant (*F*_1,18_ = 1.97, *p* = 0.18). On average, the motor preparatory activity encapsulated in the second time window (correct: 310-410 ms, error: 316-416 ms) was conspicuously linked to the response to the switch. The comparison of the LRPs generated with each correction instruction revealed a significantly higher amplitude in the preparation of correct switch responses when the instruction was to refrain any subsequent correction (*F*_1,18_ = 14.67, *p* = 0.001, *η*_*p*_^2^ = 0.45). As expected, motor preparatory activity in this second time window was larger for correct than for erroneous responses (*F*_1,18_ = 63.62, *p* = 2.6e-7, *η*_*p*_^2^ = 0.79). whereas the interaction between performance and correction instruction was again non-significant (*F*_1,18_ = 0.05, *p* = 0.82). Interestingly, motor preparatory activity of erroneous responses to switch in the second time window reflected a marginal decrease when participants were instructed to withhold the correction (*t*_(18)_ = -2.036, *p* = 0.057, *d* = -0.5), likely obeying to the fact that error corrections were not effectively suppressed on time.

**Fig. 7.**
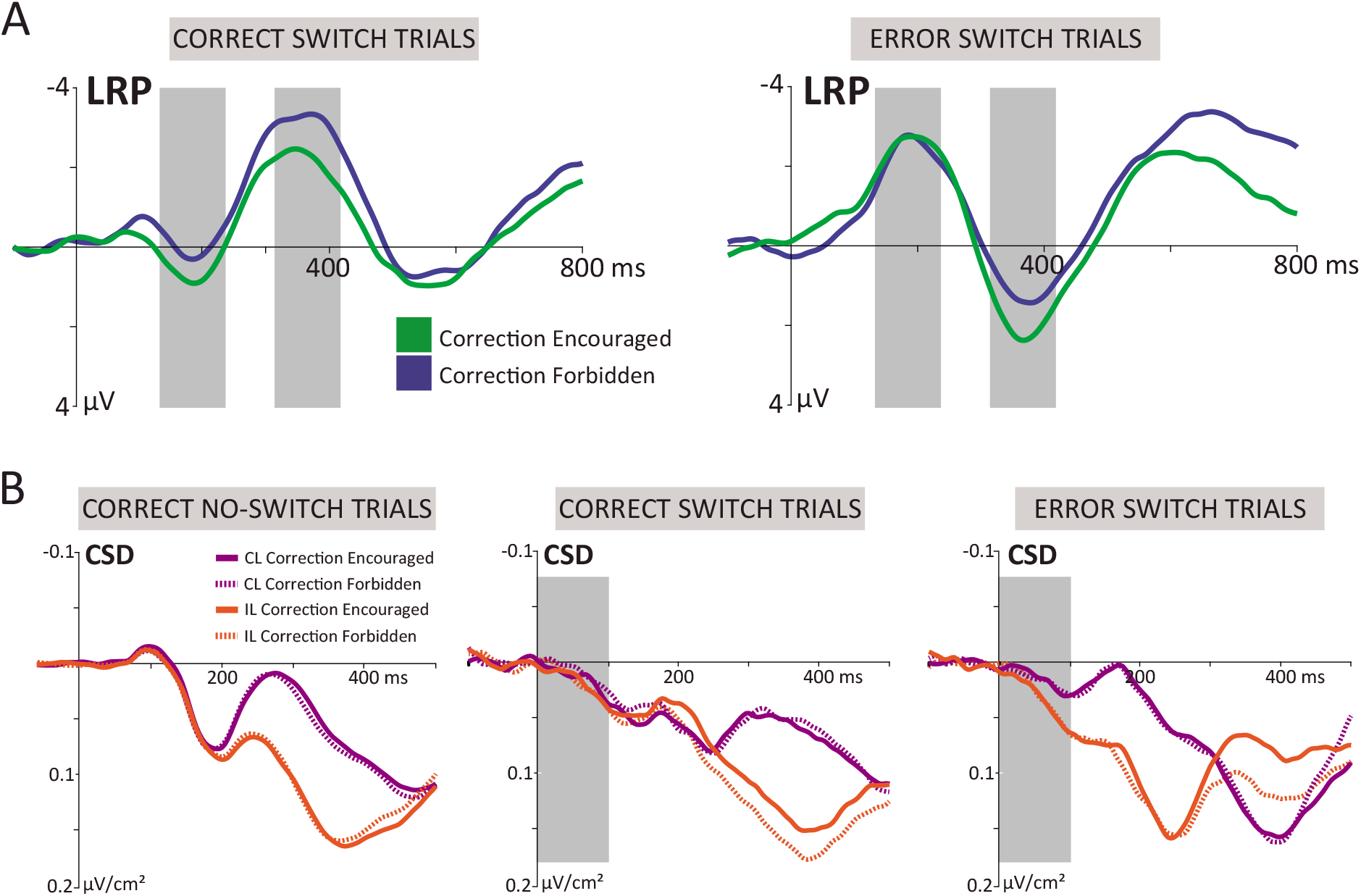
**(A)** Stimulus-locked LRPs in correct (left) and erroneous (right) switch trials for Encouraged and Forbidden correction conditions. The gray boxes indicate the analyzed time windows associated to the response preparation and to the processing of the switch. **(B)** Grand average stimulus-locked CSD waveforms of the contralateral (purple) and ipsilateral (orange) motor inhibitory activity separately for correct no-switch, correct switch and error switch trials. Solid lines represented the Encouraged correction condition whereas dashed lines correspond to trials in which the correction had to be withheld.

Lastly, the use of CSD estimates –with higher spatial resolution than ERPs– allowed us to separately quantify the contralateral and ipsilateral electrophysiological correlates associated to motor inhibitory processes with the two correction instructions. We found a positive deflection of the CSD, suggestive of motor inhibition, 100 ms after the onset of the initial stimulus. The statistical analyses in the 0-100 ms time window revealed a main effect of hemispheric laterality (*F*_1,18_ = 30.41, *p* = 3.1e-5, *η*_*p*_^2^ = 0.63), and a significant Laterality x Performance interaction (*F*_1,18_ = 11.77, *p* = 0.003, *η*_*p*_^2^ = 0.4). When decomposing the interaction, we observed that correct and erroneous responses to a switch exhibited a similar level of motor inhibitory activity in the contralateral hemisphere (*t*_(18)_ = -0.48, *p* = 0.64). Nonetheless, motor inhibitory activity in the ipsilateral hemisphere was significantly higher in correct switch trials (*t*_(18)_ = -2.94, *p* = 0.009, *d* = -0.6). These results were consistent under the two degrees of inhibitory control, and suggest that decreasing corticomotor excitability from the two hemispheres, not only the contralateral, would be required before the execution of a successful response to the switch.

To avoid a spurious baseline bias, we analyzed the CSD deflection in the same time window by calculating the mean amplitude at the beginning (0-20 ms) and at the end (74-94 ms) of the slope, and computing a linear regression for each participant and correction instruction. The baseline-free mean amplitude values were submitted to an ANOVA using the same factors and the results reproduced the significant main effect of laterality (*F*_1,18_ = 11.09, *p* = 0.004, *η*_*p*_^2^ = 0.38) and a Laterality x Performance interaction (*F*_1,18_ = 4.83, *p* = 0.04, *η*_*p*_^2^ = 0.21). Again, the inhibitory motor activity observed in correct and error switch trials only differed in the ipsilateral side (*t*_(18)_ = 2.05, *p* = 0.05, *d* = 0.44).

## DISCUSSION

In this study, we examined behavioral and electrophysiological markers of proactive control while participants performed a modified version of the Eriksen flanker task that allowed us to elicit both automatic and endogenously prepared suppression of responses. Restraining or encouraging the correction of errors did not affect the time course of the behavioral and neural correlates associated to reactive inhibition. In contrast, we found a decrease of corticomotor excitability in both hemispheres when participants effectively withheld the correction of an error, indicating a modulation of the inhibitory strength driven by correction instruction. These findings provide the first behavioral and electrophysiological conclusive evidence of a global proactive inhibitory mechanism governing the commission and amendment of incorrect responses.

Reactive inhibition is typically investigated with tasks including a condition in which a prepotent response tendency needs to be stopped (Friedman and Miyake, 2004). As a result, the use of the stop signal reaction time (SSRT) as an objective quantitative estimate of the time needed to abort an already-initiated response has been broadly supported (Aron and Poldrack, 2006; Verbruggen and Logan, 2008b). On average, SwSRT –the equivalent measure to SSRT in this study– was practically indistinguishable when participants were instructed to correct or to withhold an error just committed. Moreover, the trial-by-trial tracking exhibited a robust similarity of the SwSD under the two degrees of inhibitory control, stabilizing slightly below 100 ms the delay at which a response could still be effectively suppressed. These findings suggest that the internal speed of reactive stopping was not modulated by changes in the degree of inhibitory control. We also observed that initiating correct responses to the switch took significantly longer compared to a previous erroneous trial, reflecting careful response strategies reminiscent of the ‘switch cost’ that underlies task-set reconfiguration processes (Monsell, 2003). Contrarily, two consecutive erroneous responses to the switch showed comparable RTs. Our data thus corroborates previous evidence for flexible trial-by-trial behavioral adaptations after failed inhibitions (Verbruggen and Logan, 2008a).

We provide substantial electrophysiological evidence to warrant the validity of the two degrees of inhibitory control and rule out possible confounding factors that could mislead the interpretation of the results. One example is the replication, with the two correction instructions, of the well-established conflict-related N2 component. We reproduced the greater amplitude for incongruent –relative to congruent– trials, interpreted as evidence for the inhibition of the distracting flankers to enable the execution of the correct response (Bartholow et al., 2005; Folstein and Van Petten, 2008). Besides, the ERN-Pe compound was identified immediately after the commission of an error, as the output of an evaluative system engaged in monitoring motor conflict (Botvinick et al., 2001; Rodrıguez-Fornells et al., 2002). Remarkably, both the amplitude and the latency of the ERN elicited when error correction was forbidden were virtually identical to that of the Correction Encouraged condition, indicating that failing to halt the correction was also computed as an error-related process. The larger error-related positivity of the Pe component elicited when error correction was forbidden could be reminiscent of a stronger subjective assessment of the error, likely due to the fact that errors cannot be amended. And vice versa, being aware of the possibility to rectify an error likely reduced error significance, depicting a shorter Pe amplitude. Collectively, our results are in agreement with the view that Pe cannot be a correlate of error correction, since it was present in both corrected and uncorrected error trials (Falkenstein et al., 1996; Falkenstein et al., 2000).

The amplitude of the frontal N2 component when the response was effectively inhibited was modulated by the two different degrees of inhibitory control. Also, the larger N2 amplitude observed in unsuccessful inhibitions dovetails well with previous reports showing an increase of the N2 component in failed stop trials (Dimoska et al., 2006; Greenhouse and Wessel, 2013). One might argue though that the N2 frontal differences reflect an overlap with conflict resolution processes exclusively when a correction is encouraged (Kramer et al., 2011). This is in agreement with the view that N2 is more likely to reflect attentional processes related to the stop signal, rather than the success of an inhibitory process (Schroger, 1993; Senderecka et al., 2012).

A large body of literature supports the relationship between the frontocentral P3 waveform and successful response inhibition (Kok et al., 2004; Ramautar et al., 2004; Schmajuk et al., 2006; Wessel and Aron, 2013). Nonetheless, there is still a lack of consensus on the specific neural process that P3 subserves. We observed a larger P3 amplitude in posterior regions when participants were encouraged to correct their errors, which could tentatively reinforce its association with proactive inhibitory processes sustained over time. However, several studies refuse to link the P3 with response inhibition per se, but rather with a post hoc evaluation of the performance (Kok et al., 2004; Huster et al., 2013). This interpretation is supported by the fact that P3 peaks too late relative to the SSRT (Dimoska et al., 2003), albeit other studies have suggested P3 onset latency as a reliable neural marker of response inhibition (Wessel and Aron, 2015).

Oscillations in the beta band are known to show a prominent event-related desynchronization during the preparation and execution of manual responses (Pfurtscheller and Lopes da Silva, 1999; Hari, 2006). The absence of a concurrent decrease of the motor-related beta power when responses were not effectively withheld provides one more argument to cast doubts on the role of P3 as an index of successful response inhibition. We instead observed a frontocentral rebound of beta power starting 500 ms after the switch when participants failed to inhibit an error correction. This enhanced beta synchronization was also found to start nearly 300 ms earlier when corrections were forbidden. It is tempting to speculate that these results reflect a correlate of proactive inhibitory control triggered after the commission of an error, in line with previous studies showing a frontal beta increase associated to post-error slowing (Marco-Pallares et al., 2008).

The analysis of the LRP demonstrated that response preparation in successful responses to switch trials was inhibited as early as 150 ms after the stimulus onset, and that this pattern was consistent under the two degrees of inhibitory control. The larger motor preparatory activity observed when error correction was successfully withheld is expected, since there is no correction mechanism to be implemented and only one response must be produced. Conversely, previous studies had already shown that, when error correction was encouraged, LRPs immediately displayed a shorter lateralization interval (De Jong et al., 1995; Rodrıguez-Fornells et al., 2002). Consistent with TMS studies reporting reduced corticospinal excitability 150 ms after a stop signal (Coxon et al., 2006; van den Wildenberg et al., 2010; Cai et al., 2012), the CSD estimates reflected at 100 ms both contralateral and ipsilateral decreases of corticomotor excitability in correct responses to switch. This inhibition latency is also coincident with the estimation of around 150 ms from partial response electromyographic data (Raud and Huster, 2017). However, only the contralateral motor cortex exhibited substantial inhibitory activity when participants failed to withhold the correction. This finding converges well with the weaker motor inhibitory activity in the ipsilateral primary motor cortex observed on incompatible trials (Burle et al., 2016). Our results suggest that early bilateral changes of corticomotor excitability might be a crucial component of the cascade of processes that ultimately result in a successful response suppression.

Under the premises of the DMC framework, the proactive control mode would require the sustained maintenance of a particular type of information –the goals and rules of a task– for an optimal performance. Preserving an active representation of predefined goal-related cues has been also associated with biasing attention in favor of task-relevant information (Desimone and Duncan, 1995; Miller and Cohen, 2001), which in our study could explain the anticipation of the response to the expected upcoming switch signal. To do that, our findings indicate that generating a global –bilateral– inhibition of the motor cortex would be crucial to effectively select the desired combination of stimulus features to be mapped onto the response over other competing combinations (e.g., between executing or withholding a correction). According to this view, inhibition would occur because of response competition during action selection among conflicting task-relevant representations that correspond to the goals and the rules for achieving a behavior. Our results are consistent with the main tenet of the DMC framework, and suggest that optimizing sustained attentional monitoring would enable an adequate proactive control of response inhibition. They speak to a growing debate in cognitive control research about active mechanisms of response suppression; and motivate richer models of how to cancel a prepared or initiated action. Further research is warranted to outline putative common elements of proactive inhibition across other functional domains, such as emotional and motivational impulses.

## ACKNOWLEDGEMENTS

*TC was supported by the Ministerio de Economía, Industria y Competitividad (MEIC) (Grant number: PSI2016-79678) financed by the Agencia Estatal de Investigación (AEI) and Fondo Europeo de Desarrollo Regional (FEDER) from the European Union (UE)*.

## Notes

### Competing Interest Statement

The authors have declared no competing interest.

